# Electrogenic Dynamics of Biofilm Formation: Correlation Between Genetic Expression and Electrochemical Activity in *Bacillus subtilis*

**DOI:** 10.1101/2024.10.04.616699

**Authors:** Adel Yavarinasab, Jerry He, Abhirup Mookherjee, Nikhil Krishnanc, Luis Ruiz Pestana, Diana Fusco, Dan Bizzotto, Carolina Tropini

## Abstract

Bacterial biofilms are structured microbial communities that play a big role in diverse processes such as nutrient cycling and bacterial pathogenesis. Biofilms are known for their electron transfer properties which are essential for metabolic processes, microbial survival, and maintaining redox balance. In this study, we investigated the electrogenic properties of *Bacillus subtilis*, a bacterial producer of electron-donating biofilms. Interdigitated gold electrodes were utilized to continuously measure the electrochemical activity of biofilm-forming *B. subtilis* cells as well as genetic mutants unable to create them (biofilm-deficient), over three days of growth. The formation of extracellular polymeric substances (EPS) and filamentous appendages was monitored via scanning electron microscopy (SEM). Chronoamperometry was used to assess electrochemical activity, which showed fluctuations in electrical current at specific time points in biofilm-forming cells. In contrast, biofilm-deficient cells showed no corresponding changes in current. Cyclic voltammetry (CV) revealed significant differences between the voltammograms of biofilm-forming and biofilm-deficient cells that were hypothesized to be a result of the reduction of secreted flavodoxin only in biofilm-forming cells. Electrochemical impedance spectroscopy (EIS) was also performed at various intervals and analyzed using an equivalent circuit model. We identified the presence of a charge transfer resistance (R_ct_) exclusively in biofilm-forming cells which correlated to the time of increased electrochemical activity measured using choronoamperometry. Finally, through confocal microscopy, we found that the expression of a gene involved in biofilm matrix formation, *tasA*, was correlated with the time where electrochemical charge transfer was measured. Altogether, these results indicate that electrochemical activity is primarily present in biofilm-forming cells rather than in biofilm-deficient mutants. By combining electrochemical and microscopic methods, a methodology was developed to continuously monitor the stages of biofilm formation through measurement of electrochemical activity, substantiating a correlation between the expression of biofilm genes and their electrochemical or redox activities. These data show that electrochemical activities within biofilms vary over time and there is a temporal relationship between these processes and the expression of genes responsible for biofilm development.

## 1. Introduction

Bacterial biofilms are surface-associated communities of microorganisms enclosed in a self-produced extracellular polymeric matrix [1]; they were historically considered detrimental structures, impeding flow in tubes [2] or causing antibiotic-resistant infections [3]. However, in recent years, biofilms are increasingly appreciated as valuable biological ecosystems with the ability to degrade organic matters through a series of redox reactions [4]. This shift in perception has led to an increased focus towards utilizing biofilms in various biochemical processes, including microbial fuel cells as living catalysts [5], redox signaling modulation in the gastrointestinal tract [6], wastewater treatment to degrade chemicals [7], and bioremediation in agricultural applications to break down pollutants [8], among others.

A particular structure of interest within biofilms is the extracellular polymeric substance (EPS) matrix, which provides a protective environment and enhances the survival of the bacterial community under adverse conditions [9]. Within this structured environment, bacteria engage in intricate interactions, including those mediated by electron transfer, essential for producing the primary energy source for cells, ATP [10]. The process relies on key cellular electron carriers that move electrons through various cellular redox reactions (such as acidogenic fermentation [11], sulfate reduction [12], or methanogenesis [13]), driving energy production and sustaining redox balance. Mediated electron transfer is conducted by the secretion of redox-active molecules including flavins (e.g., flavin mononucleotide in *Shewanella* spp. [14]), phenazines (e.g., phenazine-1-carboxylic acid in *Pseudomonas* spp. [15]), and quinones (such as 2-Amino-3-dicarboxy1,4-naphthoquinone in *Lactococcus lactis* [16] or 2,6-Di-*tert*-butyl-1,4-benzoquinone in *Klebsiella pneumonia* [17]). Due to the reversible nature of the redox reactions, the mediating species can participate in multiple redox cycles; therefore, even small amounts of such molecules can facilitate electron transfer. Understanding the electron transfer mechanisms is critical for optimizing applications such as microbial fuel cells or electrolysis cells [18], where efficient electron transfer improves the performance and stability of the system. However, measuring electron transfer over time and space within biofilms is still a challenge.

The heterogeneity of microbial biofilms, characterized by non-uniform structural and biochemical composition within the EPS matrix, results in significant difficulties in studying their growth and related metabolites, including electron transfer and redox activity [19,20]. Traditional approaches for monitoring biofilm growth and metabolite dynamics such as multiwell plates and commercialized reactors (such as Robbin’s device [21] and Calgary device [22]) are limited to endpoint analyses, compromising biofilm integrity and altering cellular phenotypes [23]. Furthermore, chemical treatments, such as staining with dyes like crystal violet [24], may introduce artifacts by altering the biochemical and biophysical properties of bacterial cells [25]. These challenges emphasize the need for developing less invasive approaches to probe biofilm dynamics with precision. Electrochemical sensors have been utilized for the characterization and detection of metabolites within biofilms and offer a significant advancement (e.g., real-time and label-free detection) over conventional laboratory methods [26–29]. Such methods rely on the change in electrochemical parameters (e.g., conductance [30], impedance [31], or capacitance [32]) between two electrodes that are exposed to bacterial biofilms. Despite their versatility, electrochemical measurements can be challenging to perform in complex systems. Specifically, they do not provide information about the underlying mechanisms or the potential gene expression dynamics within biofilms over time and space.

In this study, we examine the electrogenic properties exhibited by the model biofilm former *Bacillus subtilis* and the temporal variations in electrochemical activities (current and impedance values) using a combination of continuous, non-invasive, and non-destructive methodologies. We employ the combination of electrochemical (chronoamperometry, CV, and EIS) and microscopy methods (scanning electron microscopy and confocal microscopy), to analyze the electrochemical activity and its time dependency for *Bacillus subtilis*, a bacterium that produces electron-donating biofilms. The formation of biofilm on the gold electrode and the consequent filamentous appendages within the matrix was initially confirmed using SEM. Then two strains of *B. subtilis* (biofilm-forming and biofilm-deficient) were grown on interdigitated gold electrodes and subjected to electrochemical analysis (chronoamperometry, CV, and EIS). Notably, in all electrochemical measurements, only the biofilm-forming strains exhibited a discernible change in current and charge transfer resistance, which indicates measurable redox activity. By utilizing fluorescent reporters of biofilm gene expression in *B. subtilis*, the spatial configuration of bacterial aggregates and their gene expressions were analyzed through confocal microscopy. It was shown that expression patterns of genes associated with biofilm formation occurred at the same time points that the electrochemical activities were observed. Collectively, these findings suggest that the electrochemical activity exhibits temporal variability, with a distinct relationship between these activities and the expression of genes responsible for biofilm development. These temporal dynamics suggest that changes in electrochemical behavior in biofilm at specific time points are closely linked to the regulation and expression of specific biofilm-associated genes. Overall, our results highlight the complex interplay between microbial metabolic processes and biofilm gene expression over time.

## 2. Experimental section

### 2.1. Strains and culture conditions

*Bacillus subtilis* strains were initially grown in LB medium for 24 hours in an incubator (Thermo Scientific – MaxQ 6000) and then deposited on the surface of a gold electrode (details below) at 30°C and the electrochemical measurements began unless otherwise indicated. *B. subtilis* NCIB 3610 containing an integrated dual reporter cassette on motility (P*hagA*-YFP) and biofilm formation (P*tasA*-*tsr*-mCherry) was used as the *B. subtilis* forming biofilm strain [33]. *B. subtilis* with a genetically deleted *EPS* operon (*ΔEPS*) was used as a biofilm-deficient *B. subtilis* strain (Figure S1) [33].

### 2.2. Electrochemical measurements

Biofilm formation was measured on an electrochemical platform (Figure 1a). Throughout all measurements, a gold interdigitated electrode (IDE) was used (Metrohm-G-IDEAU10) featuring two terminals on a glass substrate with 250 digits (125 digits on each side) each measuring 6760 µm in length (total geometric electrode area ∼0.338 cm^2^) with the digits separated by a 10 µm gap (0.0188 cm^-1^). The cells were cultured on the electrode (Figure 1b,c). A CH Instruments CHI660D potentiostat was used. For chronoamperometry and EIS measurements, one terminal was connected to the working electrode, and the other was connected to the reference/counter electrode leads from the potentiostat. Chronoamperometry was performed with an initial potential difference of 10 mV between the two interdigitated gold electrodes with measurements taken at intervals of 0.8 seconds over 48 hours. For the CV measurements, an Ag/AgCl wire (Antylia Scientific) served as the reference electrode while a stainless-steel wire (with a diameter of 0.5 mm with 3 loops of 4 mm in diameter) was utilized as the counter electrode. Both gold leads were connected together and made the working electrode. CVs were measured from -0.5 to + 0.5 V vs Ag/AgCl at a scan rate of 0.1 V/s. EIS was measured between the two interdigitated electrodes in a 2-electrode arrangement (no reference electrode was used) at a DC bias potential of 0 mV with a 50 mV RMS perturbation over a frequency range of 100 kHz to 0.1 Hz, selected to minimize perturbations to the sensing layer and optimize the signal-to-noise ratio [34]. A quiet time of 2 s was used at the beginning of the experiment. To minimize the time of the experiment, frequencies below 0.1 Hz were not collected, as they were strongly affected by noise (data not shown). The electrochemical measurements were repeated three times on separate IDEs for each of the three individual biological samples. The measurements were performed under identical conditions, including temperature and initial optical density, as environmental variables may influence the electrochemical dynamics of the biofilm [35]. All the presented data are the average of three biological samples unless stated otherwise.

**Figure 1.**
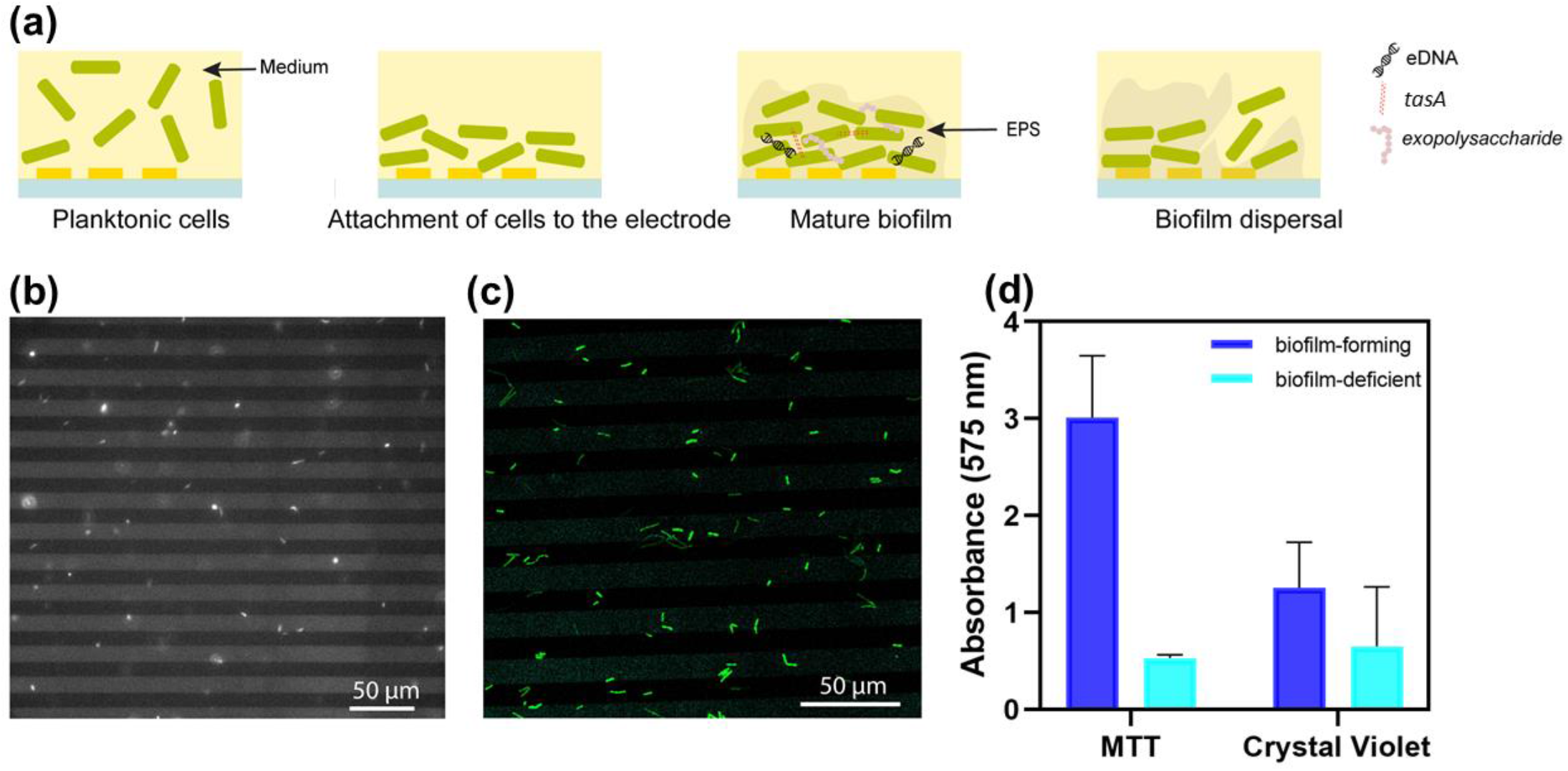
**(a)** The schematic representation of diverse stages in biofilm formation on the substrate and the presence of biofilm-related genes during the maturation phase; the culture of cells on the interdigitated gold electrodes after culturing in the incubator for 24 hours, imaging by **(b)** phase microscopy and **(c)** Fluorescent microscopy using the YFP channel (λ_ex_=514 nm) **(d)** Comparison between the absorbance of biofilm-forming and biofilm-deficient mutants using MTT and crystal violet assay after 24 hours, suggesting the formation of biofilm on the surface of the electrode.

### 2.3. Biofilm quantification assays

To assess for biofilm formation of the strains of interest, crystal violet and 3-(4,5-dimethylthiazol-2-yl)-2,5-diphenyl tetrazolium bromide (MTT) assays [36] were performed (Figure 1d). For each assay, the supernatant was carefully removed from each well following static growth at respective culture conditions in 96-well microtiter plates for 24 hours. Wells were then gently washed twice with phosphate-buffered saline (PBS -10X Solution, 1L, Fisher, Fisher BioReagents, BP3991). Absorbance readings were taken at 575 nm using a BioTek H1 Synergy plate reader to assess final biofilm formation. For crystal violet assays, 0.1% (w/v) crystal violet was added to each well and incubated for 30 minutes. Crystal violet was then removed from each well by washing twice with PBS. A 33% acetic acid solution diluted in dH_2_O was then added to the wells to resolubilize the remaining crystal violet and plates were shaken at room temperature at 550 rpm for 20 minutes. Absorbance readings were then taken at 575 nm.

For MTT assays, 0.5 mg/ml MTT diluted in PBS was added to each well and incubated at 30°C for 30 minutes. Following incubation, the MTT solution was aspirated, and dimethyl sulfoxide (DMSO) was directly added to each well to solubilize the reduced formazan crystals. Plates were then shaken at room temperature at 550 rpm for 20 minutes. Absorbance was then measured at 575 nm to assess for metabolic activity, which was used as a proxy in combination with crystal violet to quantify biofilm formation. As shown in Figure 1d, both MTT and crystal violet assay confirmed the presence of the biofilm in the biofilm-forming cells compared to the biofilm-deficient mutants.

### 2.4. Equivalent circuit modeling

The EIS was modeled using two equivalent circuits: a series RC circuit, or a Randles circuit (R in series with a parallel RC), representing an interface without and with a slow faradaic reaction, respectively. A constant phase element was used to replace the double-layer capacitance to improve the fits. A nonlinear-least-squares data fitting process using LEVM or LEVMW [37] was used to fit the data to the equivalent circuits. Choosing the appropriate equivalent circuit was accomplished using an F-test [38] as described in SI. All the equivalent circuit components were then refitted using DearEIS (https://vyrjana.github.io/DearEIS/)

### 2.5. SEM and confocal measurements

The samples were fixed with 2.5% buffered glutaraldehyde for 1 h, washed 3x with buffer, post-fixed with buffered 1% osmium tetroxide for up to 2 hours, washed 3x with distilled, deionized water, then dehydrated in an ascending ethanol series starting at 10% (20, 30, 40, 50, 60, 70, 80, 90, 95, 3×100%) on the bench (15 minutes up to 70%) and microwave (Pelco BioWave, PL3 40 sec) as shown previously [39]. The 2-hour samples were stored overnight in 70% ethanol at 4°C. After 100% ethanol, the samples were transitioned into hexamethyldisilizane (Sigma Aldrich); 50%, 75%, 100%, and 100%. The last change was heated to 65°C then decanted and allowed to evaporate at 65°C before vacuum drying overnight. The 2-hour sample was processed with ethanol in the culture dish, then the electrode was transferred into a stainless-steel critical point drying basket through completion. Due to the amount of material floating in the Petri culture dish, the 24-hour sample was processed completely in the petri dish. The electrodes were attached to 25.4 mm aluminum SEM pin stubs with double-sided carbon tape and imaged without sputter coating (Zeiss XB350). Imaging details are shown in the data zone at the bottom of the images.

Live cell confocal imaging was acquired with a Leica Stellaris 5 inverted laser scanning confocal with a Leica Application Suite X (LAS X 4.6.1), HC PL APO CS 40x/0.85 DRY objective, and HyD S detectors in xyzt mode with a time interval of 30 mins for a total time of 48 hours with a step size of 0.5 μm. YFP was excited with a 488 nm laser line and mCherry with a 561 nm laser line.

## 3. Results and discussion

### 3.1. *B. subtilis* forms robust biofilms

To investigate the morphological changes induced by biofilm formation in *B. subtilis* cells, SEM was employed. The cells were first cultured on an interdigitated electrode (Figure 2a) and then incubated statically at 30°C for about 12 hours until clusters of cells formed (Figure 2b). In the initial 2 hours, the biofilm-forming cells appeared dispersed and loosely organized, lacking the formation of the matrix (Figure 2c). However, after 24 hours of static growth, the biofilm exhibited more complex and clustered structures with a well-developed matrix and multiple layers (Figure 2d). The matrix was dense and compact, forming an interconnected network with cells tightly adhered to each other and the surface, indicating strong attachment and colonization. Micro-colonies of cells encased in EPS (consisting of polysaccharides, proteins, amyloids, lipids, and extracellular DNA (eDNA)) were also observed [9] (Figure 2e). Imaging showed cells encased in a layer of EPS, displaying a defined macromolecular “honeycomb” structure. In addition, biofilm-forming *B. subtilis* cells produced filamentous appendages, which connected the cells to each other, and possibly contributed to the integrity and stability of the biofilms (Figure 2e).

**Figure 2.**
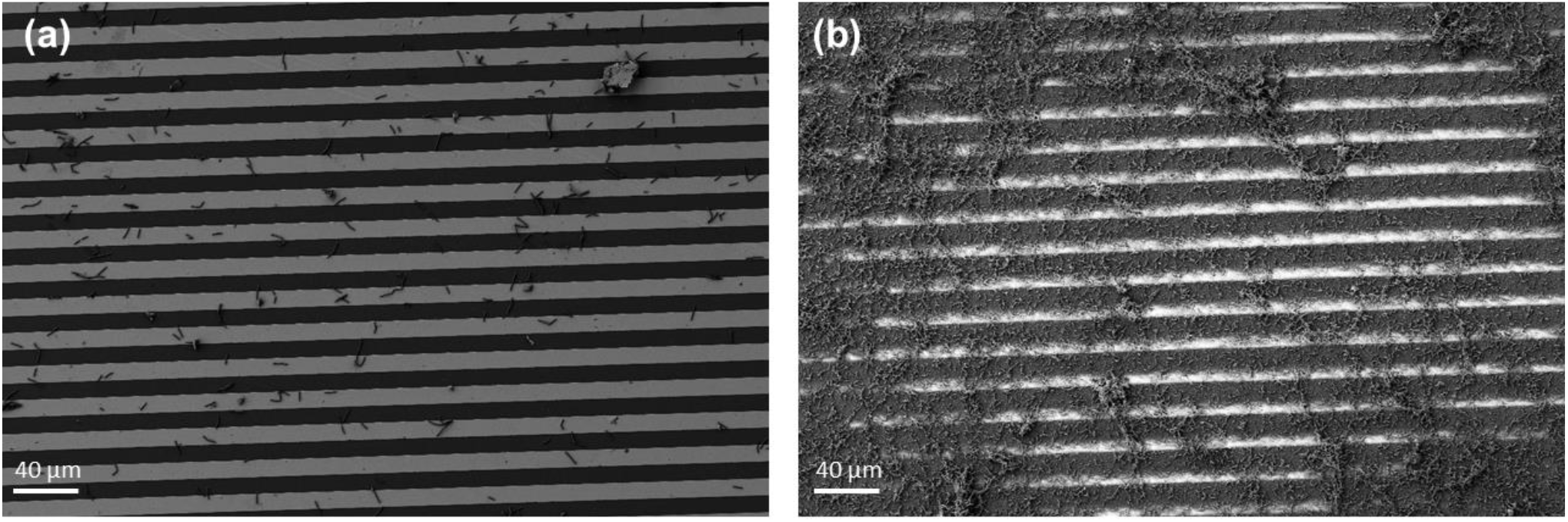
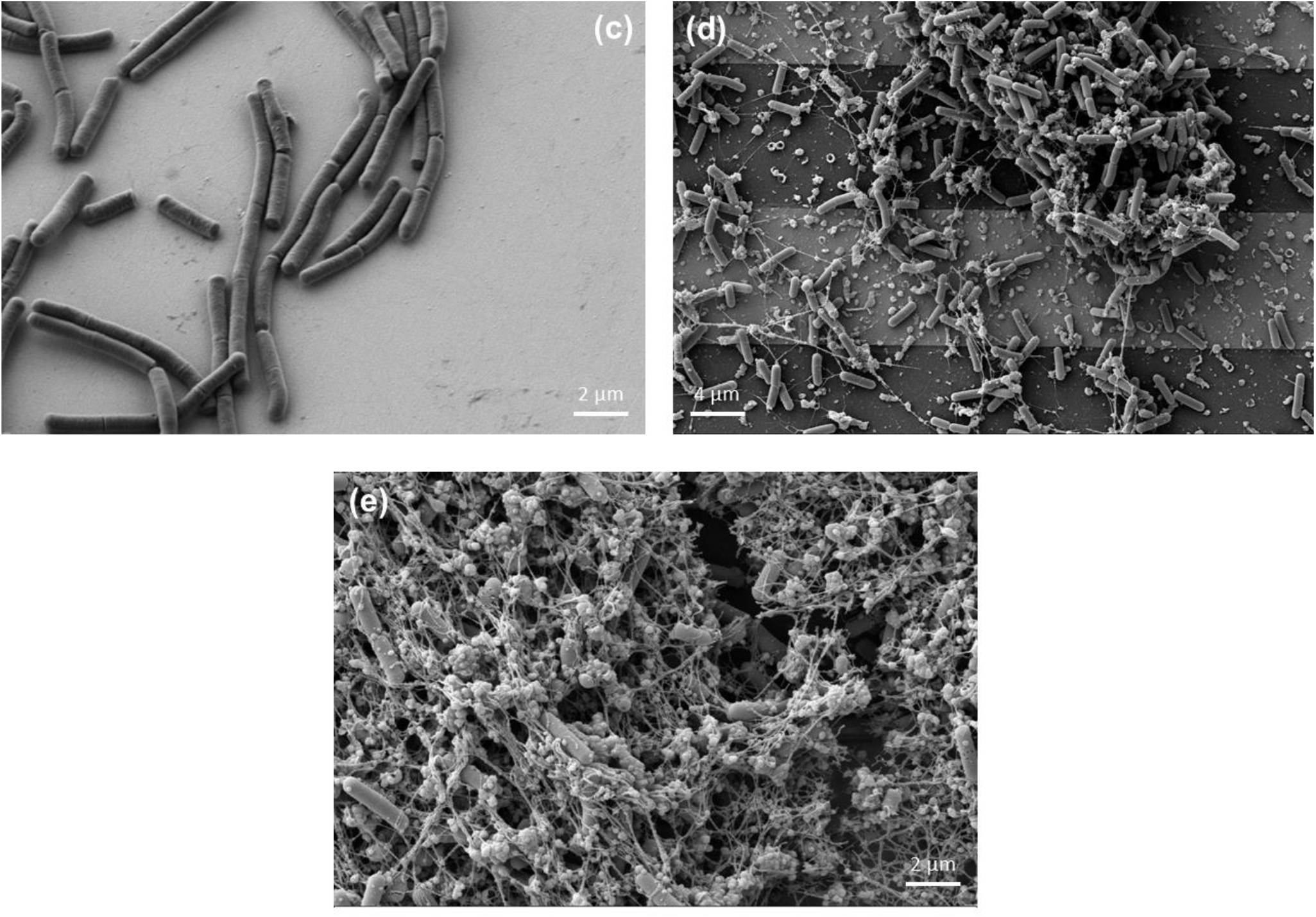
SEM images of biofilm-forming *B. subtilis* on interdigitated gold electrodes illustrate morphological changes over time: **(a)** after 2 hours and **(b)** after 24 hours of growth, highlighting the complex architecture of the biofilm. **(c)** No matrix structure is visible at the 2-hour mark. **(d)** After 24 hours, clusters of cells are attached to each other and the surface. **(e)** A detailed view of the dense EPS and the long filamentous appendages forming EPS meshes is also shown.

### 3.2. Amperometric profiling of electron transfer in *B. subtilis* shows time variability only in biofilm-forming cells

Given the robust biofilm formation observed in *B. subtilis*, the electron transfer properties were characterized via chronoamperometry during biofilm development as well as in biofilm-deficient mutants (Figure 3a). During the initial cellular attachment of the biofilm-forming cells, the current measured was notably elevated, likely due to the dense population of bacteria in close proximity to the electrode surface phase and the high respiratory activity of bacteria (with possible formation of redox species, increased consumption of oxygen, or the combination of both) in the medium. Throughout the subsequent growth phase, a progressive increase in the current was observed, indicating ongoing acclimatization and biofilm development in biofilm-forming cells [32]. Conversely, the current from the biofilm-deficient mutant remained relatively stable and close to zero over the 48-hour duration, despite the transient effects which may be due to bacterial adsorption/desorption events occurring at the electrode interface. We attribute the increase in the current measured post-10 hours to the increase of EPS thickness, alongside the upregulation of genes associated with biofilm formation (see section 3.5) [40]. Supporting these data, previous studies have shown a similar pattern of growth (e.g. [41,42]), despite differences in growth kinetics that can be influenced by factors such as medium concentration, inoculum density, electrode composition, and applied potential. Remarkably, by the 35-hour mark (± 4 hours for biological replicates), the amperometric signal in biofilm-forming cells had reverted to its initial level and the amperometric profile went back to the baseline. Figure 3b shows the CVs recorded at the electrode interface after 24 hours, both in the presence and absence of biofilm. Notably, the biofilm-deficient cells displayed little discernible redox current. This absence suggests two possible scenarios: either a slow electron transfer reduction process in these cells, or an extremely low concentration of redox-active species within the biological sample [43]. In contrast, a higher magnitude of current (more negative) was observed in the CV of biofilm-forming cells. One possible scenario to explain this phenomenon is as follows: it is known that flavodoxins, soluble electron shuttle proteins, are upregulated in *Bacillus subtilis* biofilms, with transcription peaking between 18 and 40 hours of growth [44]. These proteins are typically reduced within a potential window of -600 to -300 mV vs Ag/AgCl and play a crucial role in redox processes [45,46]. As seen in Figure 3b, CV revealed significant redox activity in this potential range, characterized by a notable decrease in current from 0 to -0.5 V vs Ag/AgCl, with minimal current observed between 0 and 0.5 V vs Ag/AgCl that aligns with the known redox potential of flavodoxin. Complementing these findings, chronoamperometry measured at 0.01 V showed a current trend that mirrored the flavodoxin transcription profile, peaking between 18 and 32 hours. These results collectively support the hypothesis that flavodoxin or a similar redox-active molecule is secreted by the biofilm and mediates extracellular electron transfer, as evidenced by both CV and amperometry data. Further investigation is required to identify the mechanisms involved in *B. subtilis* redox processes.

**Figure 3.**
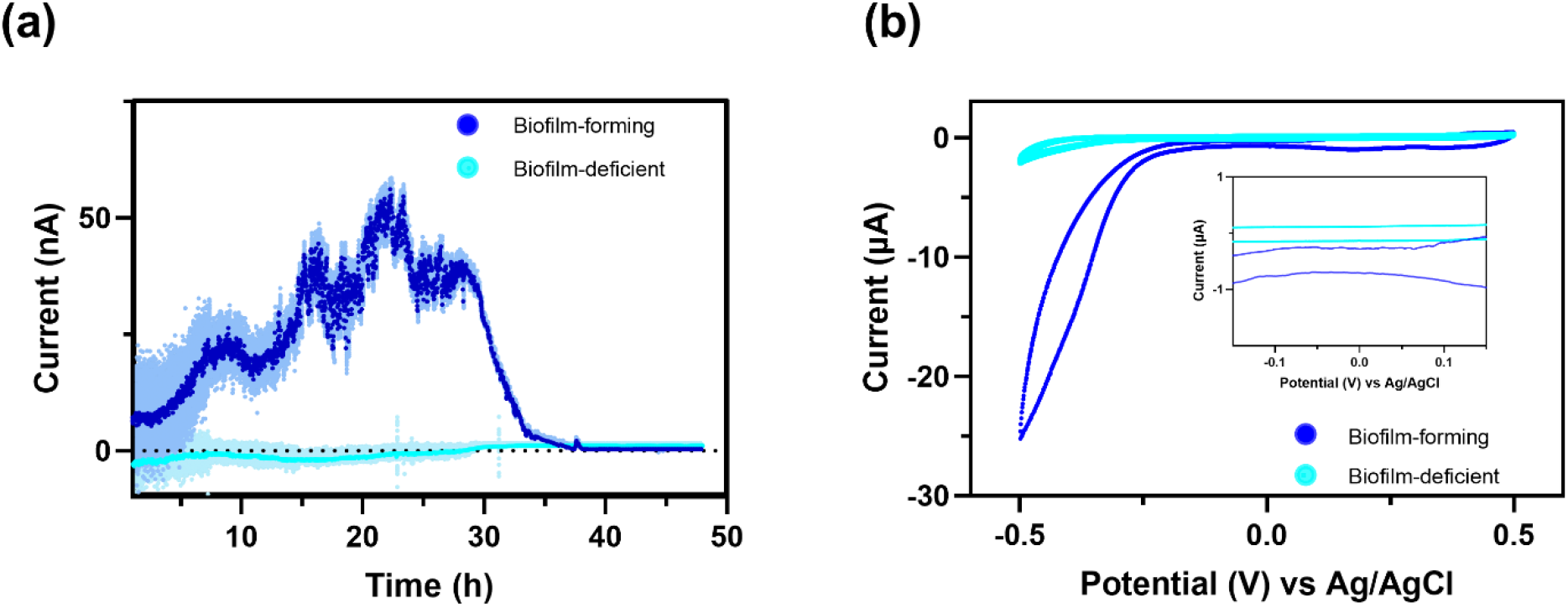
**(a)**. Chronoamperometric measurements on *B. subtilis* NCIB 3610 (biofilm-forming) and *ΔEPS* (biofilm-deficient) over a 48-hour period, starting at an initial potential of 10 mV. The measured current reflects the combined contributions of oxidation at one electrode and reduction at the other, representing the total electron transfer in the system, where the sign is not of concern. Therefore, the net current is shown as this measurement is configured as a 2-electrode measurement. **(b)** the 5^th^ cyclic voltammogram of *B. subtilis* NCIB 3610 (biofilm-forming) and *ΔEPS* (biofilm-deficient) at the scan rate of 0.1 V/s.

### 3.3. EIS analysis of *B. subtilis* dynamics shows time-dependent deviation of phase angle exclusively in biofilm-forming cells

Given the differences in electrochemical signal measured in biofilm-forming *B. subtilis* compared to biofilm-deficient, EIS was performed to provide a more detailed characterization of the interface (Figure 4). The three-dimensional Bode representations show the change in the impedance of this biological system (Figures. 4a, b, e, and f). The EIS results are the average of three biological replicates (Figure S2). Seven distinct time points were chosen for closer examination of the electrochemical response related to the generated biofilm over time. *B. subtilis* biofilm-forming cells at time points 36 hours and 51 hours were found to have phase angles at frequencies below 1 Hz deviating from 85 degrees (mostly capacitor-like in nature), decreasing to 72 degrees which are indicative of the presence of a slow faradaic reaction (Figure 4c,d). This change could also be observed when the admittance was plotted as a Nyquist plot as the beginning of the formation of a small half-circle in the low-frequency region at large Y’ (Figure S3). This suggests an increase in the amount of charge transfer compared to other time points, and the potential existence of redox molecules secreted from biofilm-forming cells. Conversely, the biofilm-deficient cells did not show a decrease in the phase of the impedance at low frequencies (Figure 4h) and therefore did not show a small semi-circle at low frequencies in the Nyquist plots (Figure S3). These results suggest the absence of redox-active species in the samples with bacteria that cannot form a biofilm. Further analysis of the impedance data can be performed by fitting equivalent circuits which would reveal quantitative changes in the electrochemical characteristics. These can then be related to the physical changes in the biofilm, given the appropriate choice of equivalent circuit.

**Figure 4.**
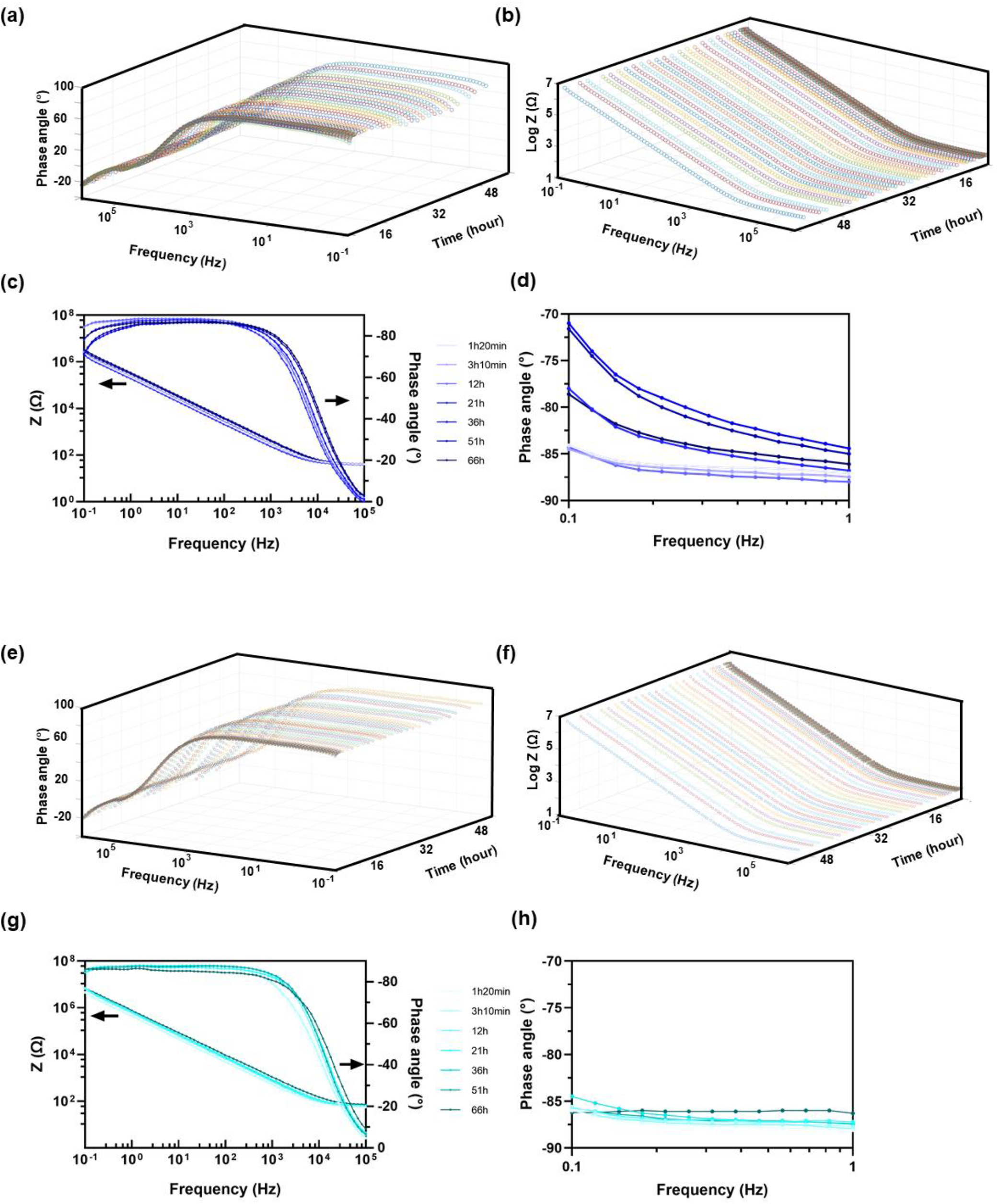
EIS representations of **(a)** the phase angle and **(b)** impedance at different time spots as a function of frequency, **(c)** the Bode diagram, and **(d)** the phase angle at the frequencies of 0.1 Hz to 1 Hz in *B. subtilis* NCIB 3610 (biofilm forming). EIS representations of **(e)** the phase angle and **(f)** impedance at different time spots as a function of frequency, **(g)** the Bode diagram, and **(h)** the phase angle at the frequencies of 0.1 Hz to 1 Hz in *B. subtilis ΔEPS* (biofilm-deficient).

### 3.4. Temporal analysis of electrochemical impedance in *B. subtilis* shows the presence of charge transfer resistance in the biofilm-forming cells

During the biofilm cultivation process, the electrochemical responses of the interdigitated electrodes were measured through continuous EIS measurements spanning a duration of 66 hours. This resulted in the acquisition of a time-response profile for each strain. An indication of the temporal change in the presence of the redox process during the growth of the cells can be estimated by plotting the impedance phase angle values at f=0.1 Hz over time (Figure 5a). As mentioned, a phase angle approaching 90 degrees (as consistently observed across all time points for biofilm-deficient mutants) indicates the interface is predominantly behaving as a capacitor. Nonetheless, departures from this capacitive behaviour at low frequencies (evident between 18 and 54 hours in biofilm-forming cells) may be attributed to the presence of faradaic or redox electrochemical processes occurring at the interface. Such deviations may arise from phenomena including adsorption dynamics, charge transfer kinetics, or diffusion constraints [47].

**Figure 5.**
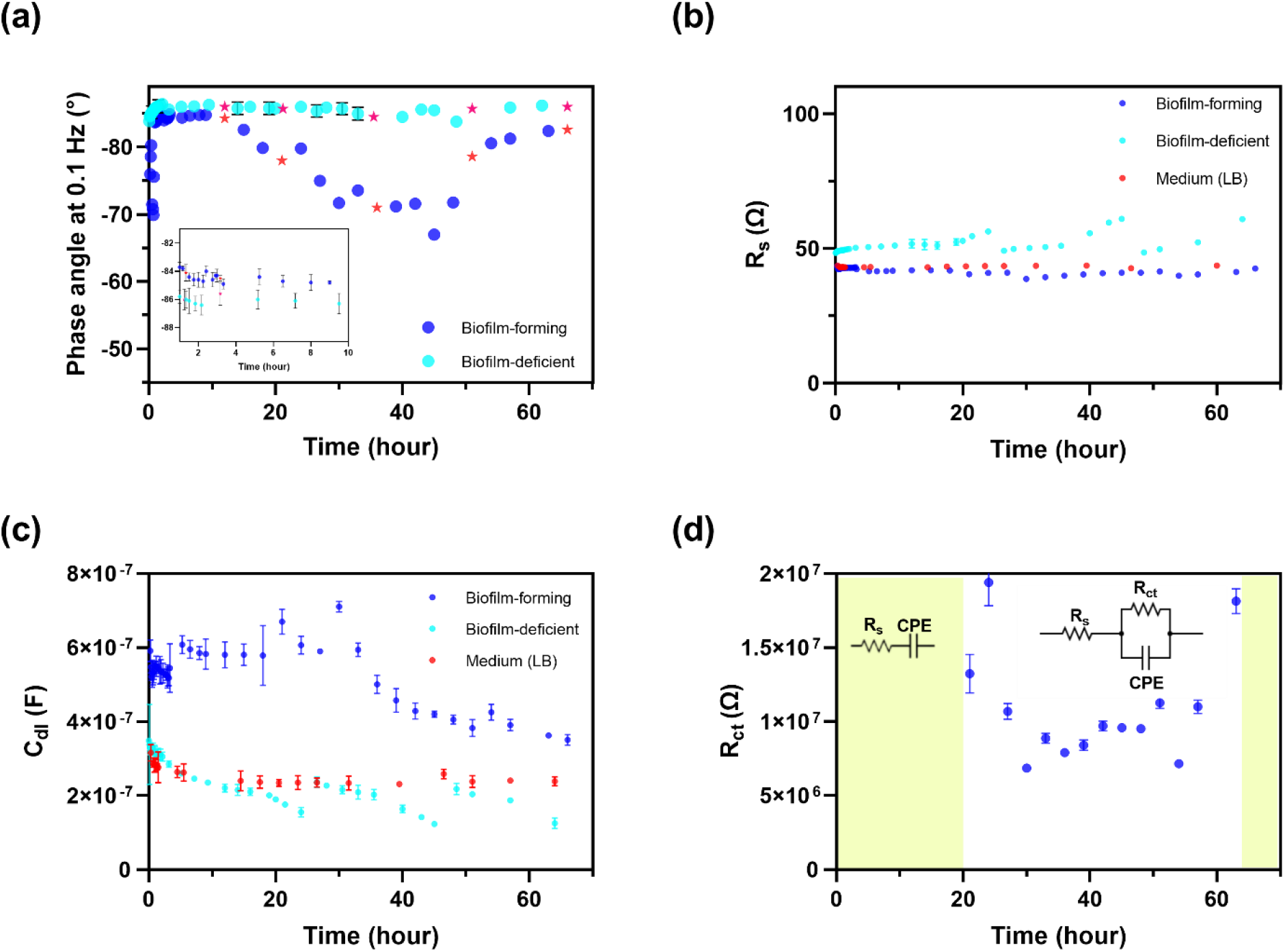
(**a)** Relative variation curves of the impedance magnitude during *B. subtilis* NCIB 3610 (biofilm-forming) culture recorded at different frequencies **(b)** The values of the phase angle at the frequency of 0.1 Hz for *B. subtilis* NCIB 3610 and *B. subtilis ΔEPS* (biofilm-deficient mutant) (Stars represent the selected data for Nyquist and Bode plots and inset is the values of phase in the first 10 hours) **(c)** Variation in double layer over time **(d)** The values of charge transfer resistance for *B. subtilis* NCIB 3610. No charge transfer was observed from the equivalent circuit in *B. subtilis ΔEPS*. (The highlighted section used a RQ circuit, and the rest was characterized by an RRQ circuit)

To interpret the physical, chemical, and biological interactions occurring at the electrode-biofilm interface, we used an equivalent circuit model to fit the EIS data [48]. This circuit model served to measure the capacitance of the interface which could be used to describe processes such as biofilm adhesion, formation, or detachment, as well as the presence of redox processes. The simplest equivalent circuit model for the strains of interest in our study was used (Figure S4). This approach avoids the possibility of overfitting the results, as multiple equivalent circuits in EIS can yield identical graphical representations; increasing the number of parameters in the equivalent circuit may enhance model fitting to empirical data [49], but decrease the usefulness of these derived parameters in terms of relating to physically relevant insights.

Specifically, the equivalent circuit adopted for the biofilm-deficient mutants across all temporal intervals comprised a simple RC circuit in series. This circuit included the resistance of solution (R_s_), representing the ohmic (bulk) resistance between electrodes in the solution, in series with the double-layer capacitance (C_dl_), arising at the electrode-solution interface due to ion solvation or adsorption of the biofilm [50]. However, due to non-ideal capacitance attributed to frequency dispersion and polarization effects, the incorporation of a constant phase element (CPE) was necessary for fitting [34]. The CPE, a versatile parameter within the equivalent circuit, has the potential to complicate the understanding of the underlying mechanistic behavior of electrochemical processes within biofilm electrodes [51] and was therefore converted into double-layer capacitance for a clearer interpretation of the electrode/electrolyte interface properties. The common formulation for CPE is 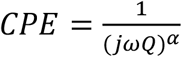, where Q and α denote the CPE parameter and exponent, respectively. The variation in CPE exponent spans from 0 to 1; approaching 1 is capacitive behavior (α = 1 denotes pure capacitance with a phase angle of 90°), while values close to 0 denote resistive behavior. In our system, we observed α values consistently ranging between 0.95 and 0.97 for both strains (Figure S5). Brug’s equation is generally used to estimate the double-layer capacitance from the equivalent circuit [52]. For a simple RQ (resistance and CPE in series) equivalent circuit

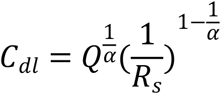

In the case of using an equivalent circuit including a charge transfer resistance (R_ct_) parallel to CPE, (RRQ) the C_dl_ can be determined using:

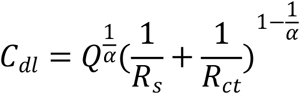

This RRQ equivalent circuit was only chosen if the addition of the R_ct_ was warranted via an F Test as discussed in the methodology and SI. In short, for all biofilm-deficient cells, the RQ circuit was employed, and no indication of charge transfer resistance was detected based on the F-test. Conversely, for biofilm-forming cells, the RQ circuit was used during the first 18 hours, followed by the RRQ circuit thereafter (Figure S4). The solution resistance remained constant throughout the experiment (Figure 5b).

Importantly, with the calculation of C_dl_, the different systems can be compared. A comparison of the C_dl_ for the electrodes in the medium alone, with biofilm-deficient cells and biofilm-forming cells is shown in Figure 5c. While the capacitance of the biofilm-deficient cell system was similar to the medium alone, the presence of a biofilm resulted in an increase in capacitance. Furthermore, the biofilm-forming system showed a larger C_dl_ from the start of the experiment, or the initiation of culturing the cells in a static mode on the electrode, following a 24-hour incubation period. In biofilm-forming cells, the dielectric constant of the EPS matrix was notably higher compared to biofilm-deficient cells or the medium, presumed to be largely due to the high content of polysaccharides within the EPS that can trap and retain water molecules as well as ions. Specifically, the polarity of polysaccharides facilitates the formation of a water-rich and ion-rich environment, which increases the dielectric constant and consequently the double layer capacitance. A slight decrease in C_dl_ over time was observed for the biofilm-forming system which was attributed to the presence of bacteria and/or EPS on the surface (Figure 5c). This phenomenon has been consistently observed across various bacterial biofilms, such as *E. coli* [53] and *P. aeruginosa* [54], and can be due to the overproduction of EPS and impaired motility while also suggesting the role of protein adsorption on electrode surfaces in modulating double-layer capacitance [55].

To investigate biofilm redox activities, the R_ct_ from the EIS analysis was analyzed over time (Figure 5d). Statistically relevant values of R_ct_ were measured solely in biofilm-forming cells after 18 hours (RRQ circuit -Figure 5d inset). Conversely, no charge transfer was observed with biofilm-deficient cells (RQ circuit). These results from the EIS analysis quantify the change in the phase at low frequencies described previously. The decrease in R_ct_ observed from 20 to 70 hours reflected the measurement of redox currents which were small. Further analysis would require identifying the redox species so an estimate of the concentration could be performed. Even so, the R_ct_ values are relatively constant over the time of 20 -70 hours, suggesting a consistent production of redox-active species which then drops off for t>70 hours. This could be interpreted as the dispersal of biofilm (Figure 1a) which can be observed as a decrease in the C_dl_ to values that are similar to those measured in the medium only.

### 3.5. Spatial dynamics through confocal microscopy show the expression of biofilm formation genes correlated with times of charge transfer

Our data identified charge transfer occurring in biofilm-forming cells at specific time points (after 18 hours). Considering the spatial heterogeneity of biofilms, we aimed to explore whether variations in the timing of gene expression in biofilm-forming cells may be linked to changes in electrochemical signals. We hypothesized that alterations in the redox activity of *B. subtilis* biofilms would be correlated with expression profiles of major biofilm genes. Among genes involved in *B. subtilis* biofilm formation, *tasA* and *hag* are of key interest, having been shown to be crucial in colony and pellicle biofilm formation and encoding flagellin, important for motility and biofilm production, respectively [51]. To investigate changes in gene expression during the period of electron transfer in biofilm-forming cells (24 to 40 hours, Figure 5d), confocal microscopy imaging was performed at intervals of 30–60 minutes on cells expressing fluorescent reporters of these genes. As depicted in Figure 6, during the initial 2 hours of culturing, when no charge transfer from the EIS data (Figure 5) occurred in the biofilm, the *tasA* gene was not yet expressed, and the *hag* gene was expressed at low levels. TasA serves as an assembly and anchoring protein for the biofilm matrix and facilitates bacterial adhesion to surfaces, thus initiating biofilm formation [56]. We found that the charge transfer resistance began to emerge in the equivalent circuit of the EIS data over time (after 24 h), at a time when the *tasA* gene also increased in expression (Figure 6). Furthermore, we observed that when the charge transfer value remained relatively stable in the equivalent circuit (between 27 and 33 hours – Figure S6), both the *tasA* and *hag* genes were expressed consistently with only minor changes, as shown in Figure S6. Notably, while biofilm-deficient cells failed to adhere to the substrate (image not provided) resulting in C_dl_ values that were not different than the measurement in media alone, the C_dl_ values in biofilm-forming cells (Figure 4c) showed the formation of substantial biofilm structures and the production of EPS (Figure 2d-e and Figure 6). Importantly, we observed the expression of *tasA* and *hag* genes only after R_ct_ was detected in the equivalent circuit from the EIS data (beyond 24 hours). These data indicate that these genes may correlate with the regulation of biofilm redox activity and the charge transfer dynamics between the electrode and the EPS matrix.

**Figure 6.**
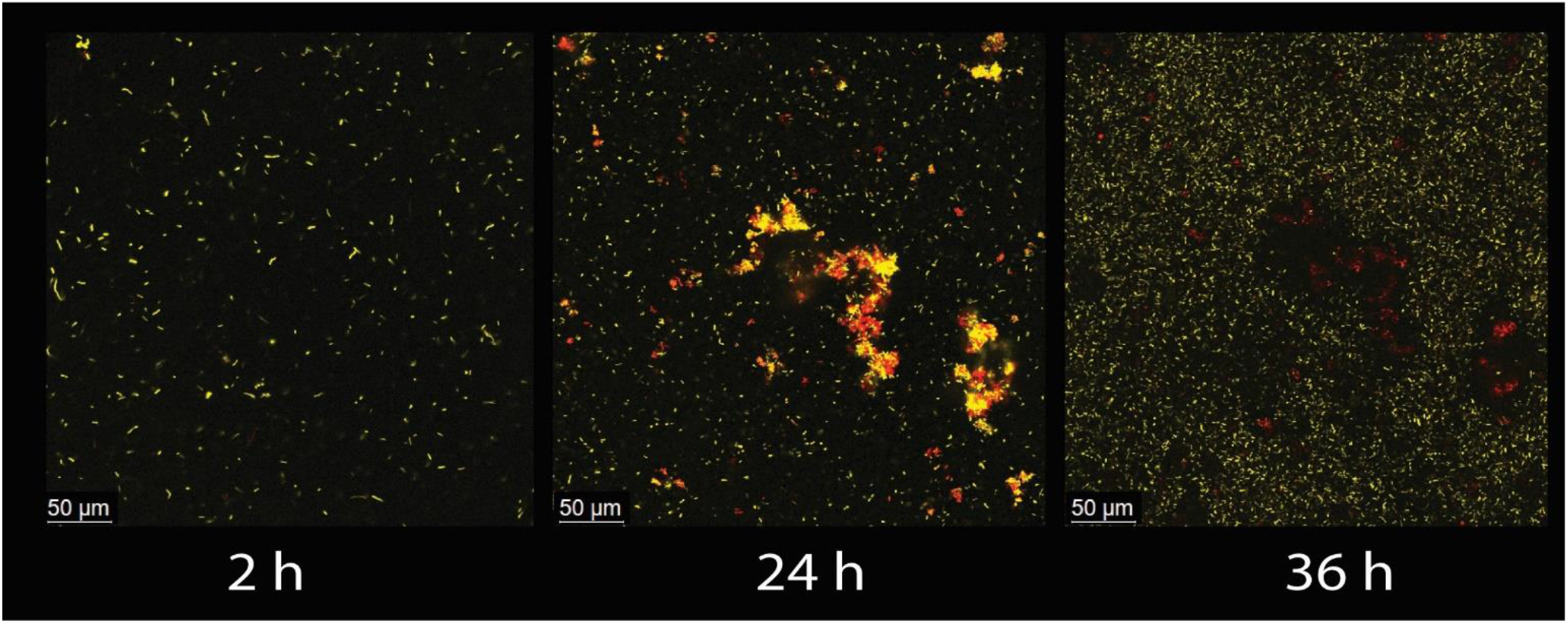
Dual channel confocal laser scanning microscopy micrograph of *B. subtilis* NCIB 3610 *sacA::Phag-yfp*-*amyE::PtasA-tsr-mcherry* biofilm different time points; No expression of *tasA* (red) was observed after 2 hours of culturing, coinciding with the absence of charge transfer resistance in EIS measurements. After 24 hours, when charge transfer appeared in the EIS circuit, both *hag* (yellow) and *tasA* were expressed. By 36 hours, with consistent charge transfer resistance in EIS measurements, *hag* and *tasA* were stably expressed at different locations on the surface. Colour allocation: Red: mCherry reporter for *tasA* gene; yellow, YFP reporter for *hag* gene

Interestingly, in other studies, it has been suggested that cells in different layers of biofilms use separate electron acceptors [57]. Specifically, upper-layer cells were found to use molecular oxygen for intracellular electron transfer, whereas lower-layer cells, encountering oxygen limitation, utilized extracellular matrix-bound iron for extracellular electron transfer. However, this proposed mechanism is unlikely to hold true in our investigation, as evidenced by the lack of redox activity observed in *B. subtilis ΔEPS*, despite exposure to oxygen in upper layers (Figure 3).

## 4. Conclusion

This study shows the intricate interplay between biofilm formation and electrochemical activity in *Bacillus subtilis*. Our data shines a light on the time dependency of electron transfer within microbial biofilms and correlates it with expressions of matrix and motility genes at different time points. By employing genetic mutants, the combination of electrochemical measurements (chronoamperometry, CV, and EIS), and microscopy (SEM and confocal microscopy), we observed that the expression of biofilm-related genes (*tasA* and *hag*) correlate with the time of high charge transfer resistance. We further showed how EIS can be used continuously to monitor bacterial biofilm and can complement other traditional electrochemical methods such as amperometry or voltammetry, where the information about the electrode/electrolyte interface is limited, or hard to interpret. Our findings underscore the importance of *tasA* in mediating biofilm adhesion and facilitating the initiation of biofilm formation, which correlates with changes in the electrochemical characterization. Importantly, our approach holds promise for future applications in optimizing systems where electron transfer can improve efficiency such as in bacterial fuel cells. Future work involving transcriptomics and the creation of genetic mutants will be required to uncover the underlying mechanisms of electron transfer in bacterial biofilms.

## Acknowledgments

The authors acknowledge that the land we performed this research on is the traditional, ancestral, and unceded territory of the xwmƏθkwƏýƏm (Musqueam) Nation. The land it is situated on has always been a place of learning for the Musqueam people, who for millennia have passed on their culture, history, and traditions from one generation to the next on this site. We encourage others to learn more about the native lands in which they live and work at https://native-land.ca/

The authors also acknowledge the UBC Bioimaging Facility (RRID: SCR_021304) along with Derrick Horne and Miki Fujita for SEM image acquisition and technical support as well as the UBC LSI Imaging Core Facility (RRID: SCR_023783) along with Guang Gao for confocal microscopy support. We also thank Camden Hunt and Curtis P. Berlinguette for the electrochemical potentiostat station. Finally, the authors thank members of the Tropini and Bizzotto labs for their support of this work.

