## Supplemental Figures for "Electrogenic Dynamics of Biofilm Formation: Correlation Between Genetic Expression and Electrochemical Activity in *Bacillus subtilis*"

**Supplementary:**

In the fitting of EIS data to an equivalent circuit, it is acceptable to incorporate an additional parameter into the model only if it yields a statistically significant reduction in the sum of squares [38] as follows:

$$F_{exp} = \frac{\frac{S_1 - S_2}{(2N-m)-(2N-m-k)}}{\frac{S_2}{2N-m-k}} = \frac{\frac{S_1 - S_2}{k}}{\frac{S_2}{2N-m-k}} = \frac{S_1^2}{S_2^2}$$

Here, $S_1$ is the sum of squares for the model containing m parameters, $S_2$ is the sum of squares for the model after adding k parameters, and 2N-m is the initial degrees of freedom. For all F-tests, the confidence level of 0.05, corresponding to probabilities of 95 % was selected.

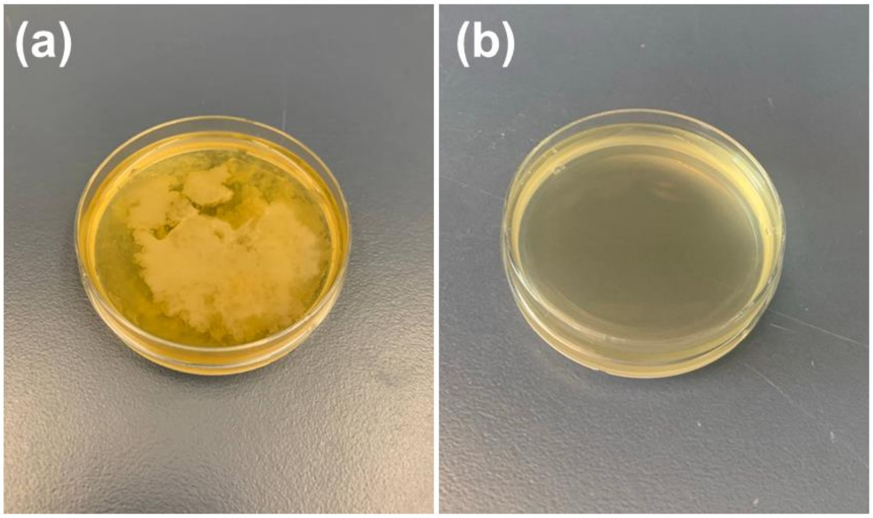


**Figure S1**. Comparison of biofilm formation in two cell samples. **(a)** *B. subtilis* NCIB 3610 shows visible cloud-like structures, indicating robust biofilm development. **(b)** *ΔEPS* mutant cells exhibit no cloud-like structures, demonstrating the absence of biofilm formation.

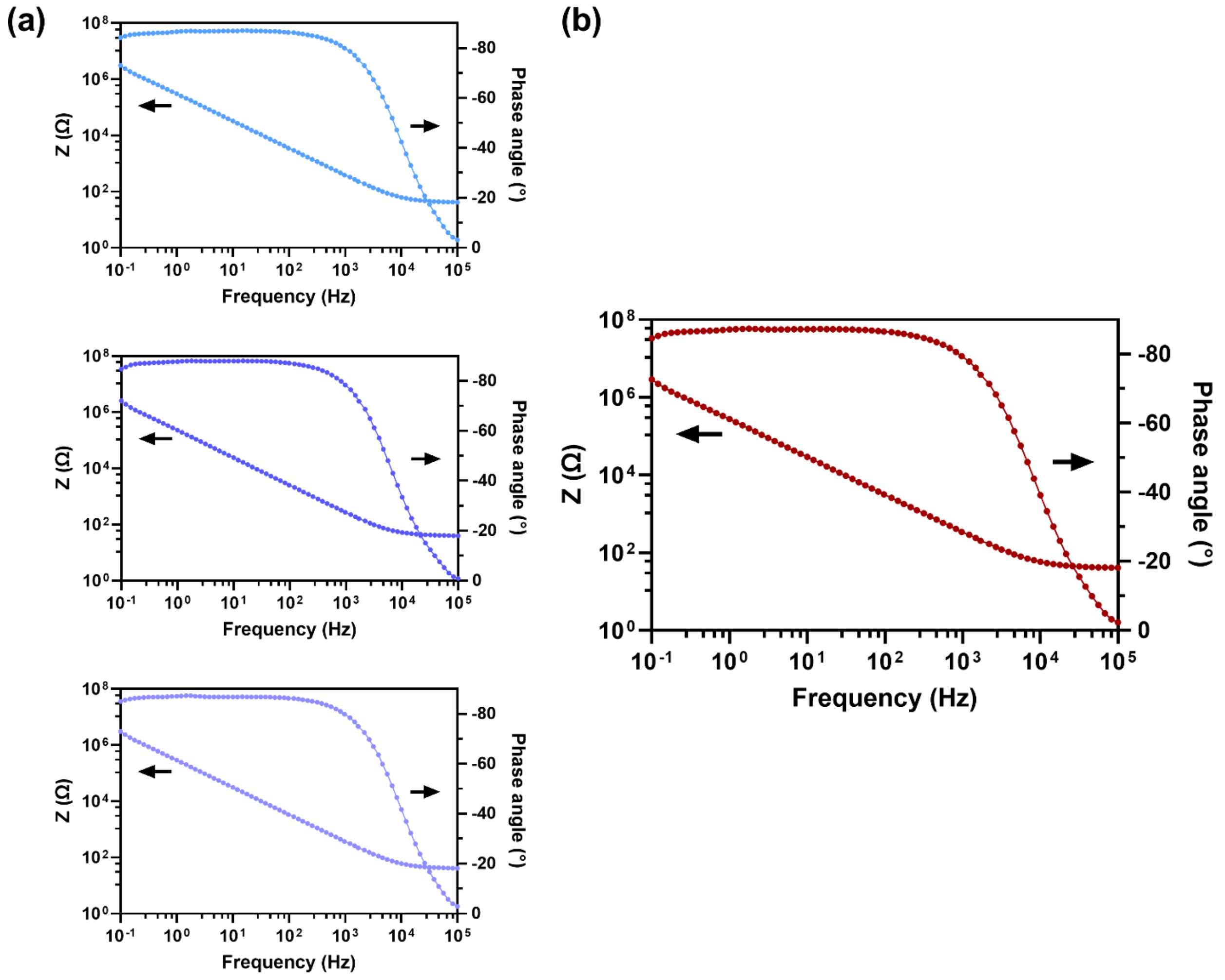


**Figure S2**. **(a)** Three biological replicates for the Bode diagram of *B. subtilis* NCIB 3610 after 15 hours **(b)** The average values of the impedance and phase angle derived from the replicates; the biological replicates were performed on three similar interdigitated electrodes

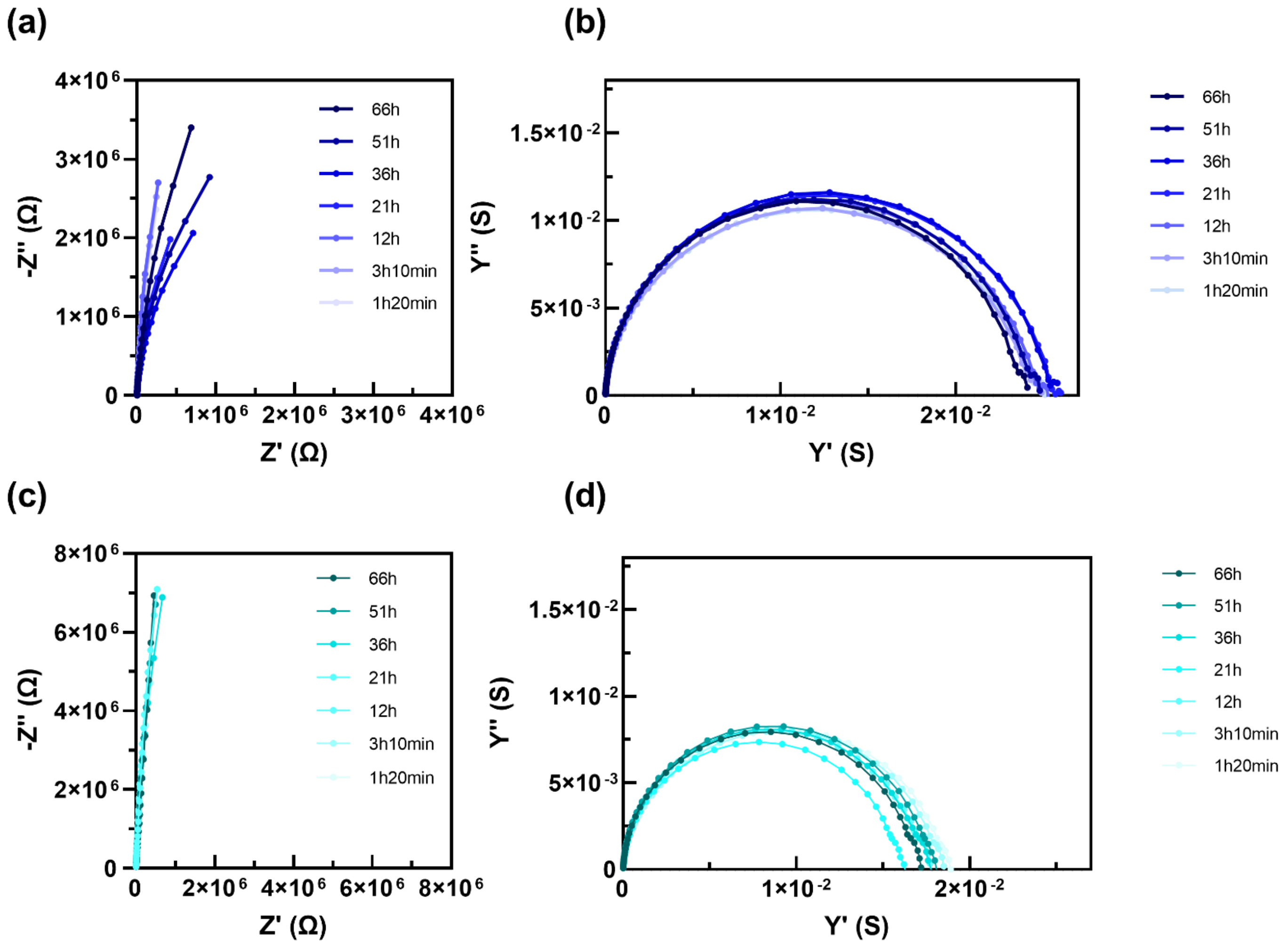


**Figure S3**. The Nyquist and admittance diagram of **(a)** and **(b)** *B. subtilis* NCIB 3610 (biofilm-forming) and **(c)** and **(d)** *B. subtilis ΔEPS* (biofilm-deficient) over different time spots

**(a)**

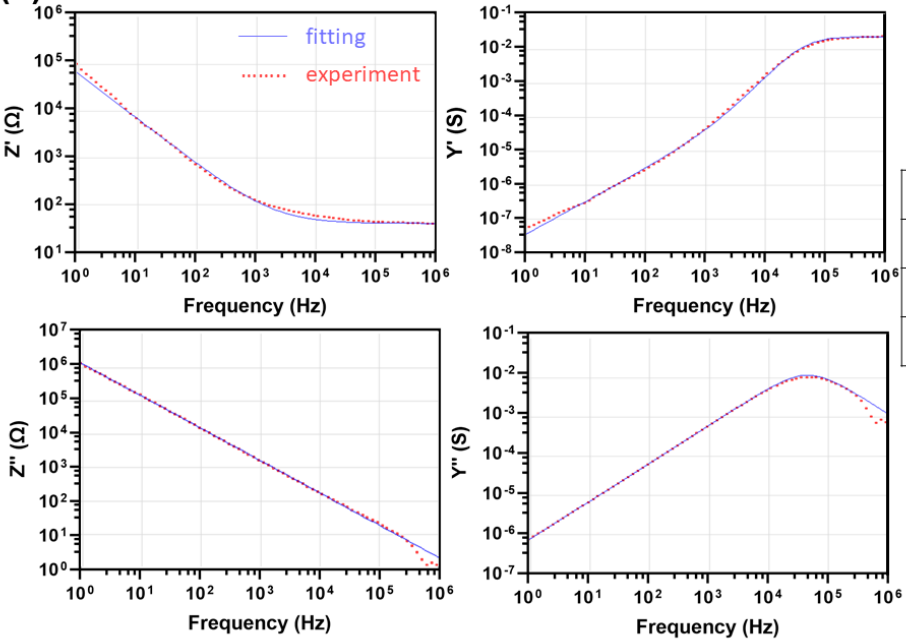


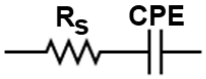


| Component | Value | RSD |
|---|---|---|
| $R_S$ | 40.2 | $2.18 \times 10^{-3}$ |
| Q | $7.67 \times 10^{-7}$ | $2.55 \times 10^{-4}$ |
| α | 0.969 | $3.20 \times 10^{-5}$ |

$\log \chi^2 = -1.84$

**(b)**

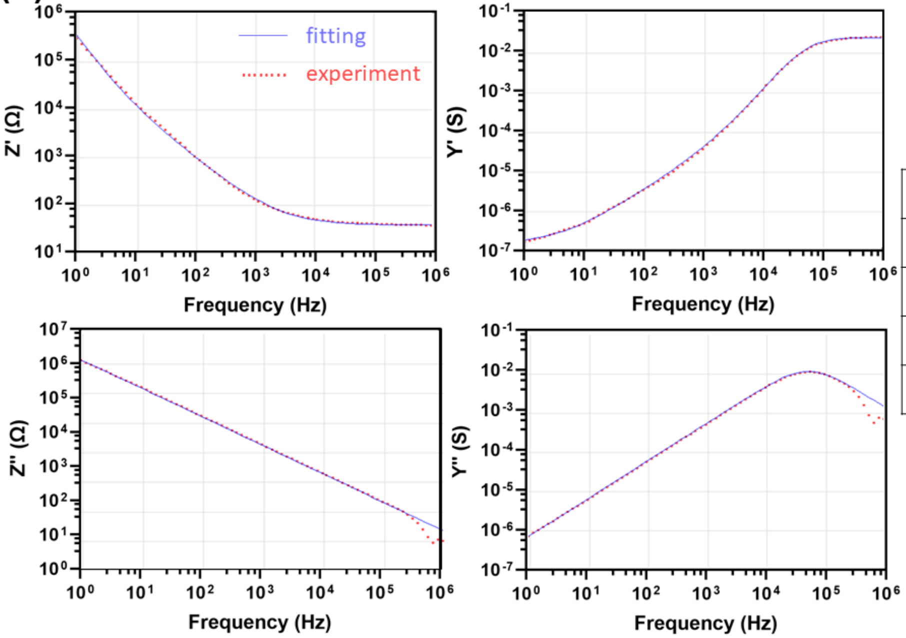


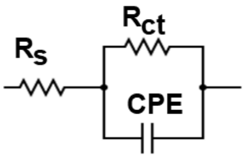


| Component | Value | RSD |
|---|---|---|
| $R_S$ | 39.93 | $5.02 \times 10^{-4}$ |
| Q | $7.40 \times 10^{-7}$ | $8.68 \times 10^{-5}$ |
| α | 0.963 | $9.49 \times 10^{-6}$ |
| $R_{ct}$ | $7.91 \times 10^{6}$ | $3.91 \times 10^{-3}$ |

$\log \chi^2 = -3.91$

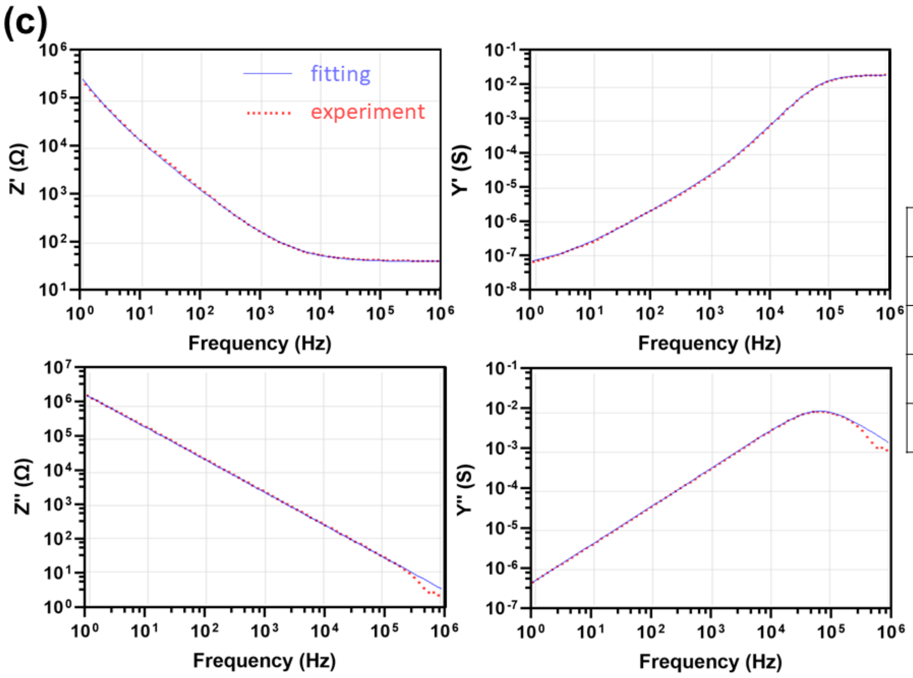


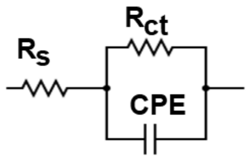


| Component | Value | RSD |
|---|---|---|
| $R_S$ | 42.61 | $9.21 \times 10^{-4}$ |
| Q | $4.86 \times 10^{-7}$ | $2.84 \times 10^{-5}$ |
| α | 0.97 | $5.44 \times 10^{-6}$ |
| $R_{ct}$ | $2.56 \times 10^{7}$ | $5.92 \times 10^{-3}$ |

$\log \chi^2 = -4.00$

**Figure S4**. Comparison between the experimental data and the fitting result from the two used equivalent circuits for real and imaginary components of impedance, and real, and imaginary components of admittance as a function of frequency and the values of the components of the equivalent circuit for **(a)** 4 hours, **(b)** 36 hours, and **(c)** 66 hours; RSD = relative standard deviation

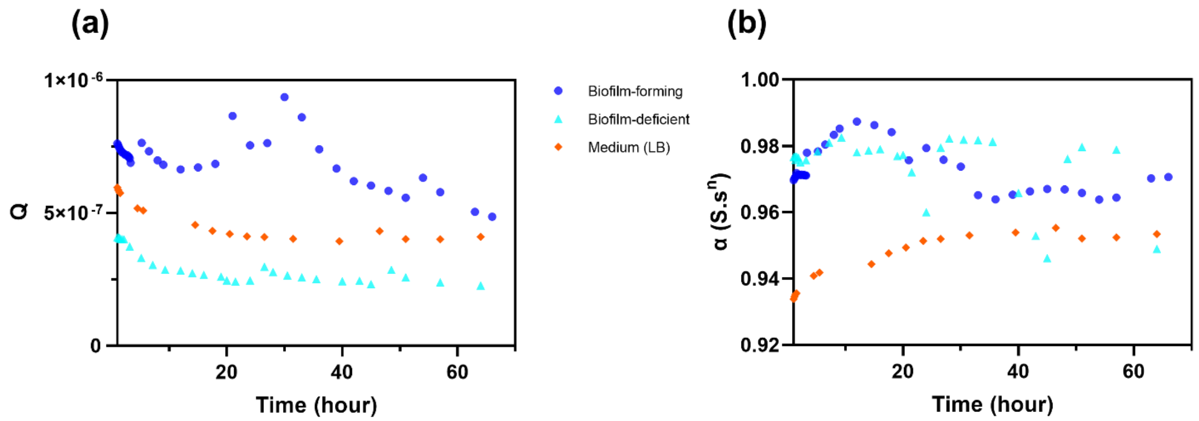

**Figure S5**. Variations in the values of CPE components **(a)** parameter and **(b)** exponent used in the equivalent circuit for the biofilm-forming, biofilm-deficient, and the medium as the control

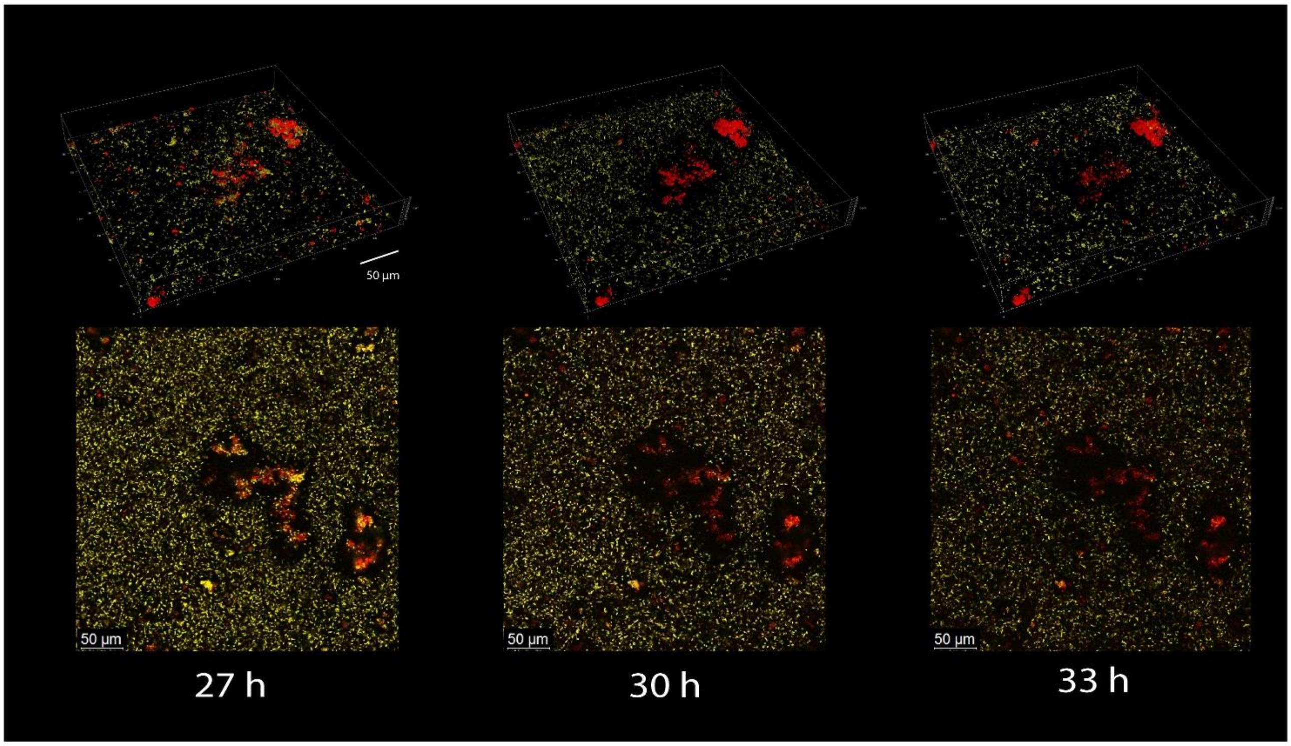


**Figure S6**. Confocal microscopy conducted between 27 and 36 hours shows minimal changes in gene expression that matches the time point where charge transfer is occurring in EIS
